# Deciphering biological evolution exploiting the topology of Protein Locality Graph

**DOI:** 10.1101/2021.06.03.446976

**Authors:** Barnali Das, Pralay Mitra

## Abstract

The conventional sequence comparison-based evolutionary studies ignore other evolutionary constraints like interaction among proteins, functions of proteins and genes etc. A lot of speculations exist in literature regarding the presence of species divergence at the level of the Protein Interaction Networks. Additionally, it has been conjectured that the intra-module connections stay conserved whereas the inter-module connections change during evolution. The most important components of the biological networks are the functional modules which are more conserved among the evolutionary closer species. Here, we demonstrate an alternative method to decipher biological evolution by exploiting the topology of a spatially localized Protein Interaction Network called Protein Locality Graph (PLG). Our lossless graph compression from PLG to a power graph called Protein Cluster Interaction Network (PCIN) results in a 90% size reduction and aids in improving computational time. Further, we exploit the topology of PCIN and demonstrate our capability of deriving the correct species tree by focusing on the cross-talk between the protein modules exclusively. Our results provide new evidence that traces of evolution are not only present at the level of the Protein-Protein Interactions, but are also very much present at the level of the inter-module interactions.

## 1 Introduction

Recently, a combination of the objectives from evolutionary biology and systems biology resulted in the birth of a new biological research domain named as the evolutionary systems biology. A majority of the works on the evolutionary systems biology is about species divergence and evolution by performing genome and proteome sequence-based analyses ^1–4^. Such conventional sequence comparison-based methods ignore evolutionary constraints like the structure and function of genes and proteins ^5^. A comparative strategy has been developed by Liang et al. for a global analysis of the Protein Interaction Networks (PINs) of different organisms ^6^. A combination of interaction topology and sequence similarity has been exploited for identifying the conserved network substructures. Despite the incompleteness of protein interaction data, a comparison of the PINs disclose species conservation at the network level ^7^. Also, the identified network substructures are found to be functionally conserved and they involve in essential cellular functions. Jin et al., simultaneously, analyzed the PINs of multiple organisms to study the evolutionary dynamics of the PINs ^8^. On the other hand, Zhong et al. perform inter-interactome mapping to gain insights into the evolution of biological networks ^9^. Overall, network comparisons aim to provide essential biological knowledge that goes beyond what is obtained from the genome ^10^.

The most common data utilized for the reconstruction of the phylogenetic trees is the sequence data which has already provided evidence for the the evolutionary mechanisms like deletion, mutation, and insertion. Over the past decade, similar to the sequence data, the amount of available protein interaction data has been steadily increasing. Despite an ample amount of PPI data now available, to the best of our knowledge, there exist few or no phylogenetic tree reconstruction methods based purely on the Protein Interaction Networks (PINs). Yet, a lot of conjectures exist in the literature regarding the presence of traces of evolution at the level of the PINs. Even though evolution is usually studied at the genomics level, a sizeable number of researchers claim and support that there must also be an obvious presence of species divergence at the level of the PINs ^6,11,12^. Again, topological analysis of biological networks which provides local and global quantification of the network structure ^13^, assists in exploring the problems relating to the evolution of life and has the potential to provide key insights on the evolution of biological functions at the systems level. Ali et al. reports that in spite of the lack of conservation of individual PPIs, the topology of PINs contains relevant information about the evolutionary processes ^7^. It has also been speculated that PIN-based phylogenetic trees may give rise to a major progressive step change in evolutionary systems biology ^14^, and biology as a whole. Nonetheless, it is true that, there already exist a large number of elegant genome sequence-based methods for constructing the species trees. And, it is also true that, the currently available incomplete and error-prone PPI datasets will allow the PIN-based tree reconstruction methods to have limited success as well as they will only span a limited species set ^7^. In contrast, PIN-based phylogenetic tree construction methods will be much of interest since they will decipher the role of evolutionary mechanisms in the biological networks. Such methods will also be useful in other domains where alternating means of generating taxonomies are not available ^7^.

Hartwell et al. mentions in ^15^ that, during evolution, the core function of a module is too robust to change. However, the properties and functions of the cell, i.e. its phenotype, change during evolution by altering the connections between different modules. Again, there exist several arguments regarding the difficulty of evolution of the proteins involved in numerous different interactions ^16^. Dense connections exist at the center of the cluster whereas sparse connections are present near the periphery of the module and in between the clusters. Hence, the hub proteins and their immediate neighbors in a cluster are engaged in massive number of interactions among each other. Thus, these proteins as well as their corresponding Protein-Protein Interactions possess very little possibility of undergoing any changes during evolution. Whereas, the proteins present in the periphery of the cluster and the inter-module/inter-cluster connections alter during evolution causing phenotypic variations in the cell.

Based on the modular cell biology principles, we propose our hypothesis that traces of evolution are not only present at the level of the PINs but are also very much present at the functional modules level that are densely connected sub-networks or clusters of the PINs. We start by constructing a protein locality graph (PLG) from the whole-cell protein interaction information of the selected organisms using an automatic network-based zoning method ^17^. We reduce the PLG to a power graph called as the Protein Cluster Interaction Network (PCIN), where each functional module of the PLG is treated as a node, and the inter-cluster/inter-module connections in PLG constitute the set of edges. We analyze the topology of the Protein Locality Graph and the cross-talk between the PLG clusters. Then, we verify their roles in offering the possibility of network phylogeny reconstruction. Additionally, we propose our hypothesis and perform statistical analyses based on simulated networks to verify the hypothesis. The evolutionary tree constructed by the PCIN topology-based analysis is highly similar to that generated by the PLG topological analysis. Nevertheless, as the problem size reduces from PLG to PCIN, so the computational time is also reduced drastically in PCIN as compared to PLG. Finally, our results demonstrate that traces of evolution are not only present at the level of the Protein Interaction Networks but are also very much present at the functional modules level which are the densely connected sub-networks or the clusters of the PINs.

## 2 Methodology

### 2.1 Hypothesis

Thorough analysis performed on the experimental Protein-Protein Interaction data in ^7^ confidently concludes the presence of traces of evolution at the level of the Protein Interaction Networks. Again, comparative analyses performed on the computationally predicted PINs in ^18^ disclose that evolution of the PINs is a major contributing factor towards the evolution of the closely related organisms. The Protein Locality Graph ^17^ is a variant of PIN, which not only includes experimental PPI information from the existing experimental PPI databases, but also maps pathway information to interaction data and incorporate them into the locality graph. Interactions are also predicted based on the protein spatial locality which are also a part of the PLG. Hence, the PLG is a mixture of experimental PPI data, pathway information, and locality-based predicted PPI data. Here, firstly, we analyze and verify that whether the topology of the Protein Locality Graph contains information about the evolutionary processes or not.

Following the modular cell biology principles, the spatially localized clusters of PLG behave as functional modules. Long back around the year 1999, it has already been indicated that the intra-module connections stay conserved whereas the inter-cluster connections change during evolution ^15^. This is mainly because, it is very difficult for the proteins engaged in numerous interactions to undergo any change since it may impact their numerous corresponding interaction partners or the enormous respective interactions. Thus, the hub proteins and their corresponding immediate neighbors which share dense connections among each other and are located at the center of a cluster will stay preserved during evolution. Whereas, the proteins located in the periphery of a cluster which possess sparse connections with other peripheral proteins of other clusters resulting in the inter-cluster connections, undergo significant change during evolution ^16^. Additionally, the most important components of the biological networks are the functional modules ^19,20^, which are more conserved among the evolutionary closer species ^5^. Hence, the functional modules play an essential role in the evolution of biological systems ^21–24^.

Research conducted in ^25^ construct an Epistasis Map (E-MAP) for the fission yeast, *Schizosaccharomyces pombe*, by systematically measuring the phenotypes associated with the pairs of mutations. E-MAP is a quantitative genetic interaction map which focus on different types of chromosome function such as DNA repair/replication, transcription regulation, etc. Comparison of the *S. pombe* E-MAP to an analogous E-MAP of the budding yeast, *Saccharomyces cerevisiae*, reveals conservation of the intra-module connections at the functional modules level, whereas, substantially differing cross-talk between the modules resulting in biological evolution ^25^. This fact has been depicted in Figure 1, a result extracted from ^25^, which has been generated by performing E-MAP comparisons of the above two mentioned organisms. Figure 1 clearly shows how the intra-module connections are conserved among the two organisms, budding and fission yeast. Additionally, we can see from Figure 1, that apart from three inter-module associations which are conserved among the two species (*SWR-C — SET1-C*, *SWR-C — SET3-C*, and *Prefoldin — SET1-C*), rest of the inter-module attachments are significantly different between the two and are thus primarily contributing towards the evolution of the two closely related species, budding yeast and fission yeast.

**Figure 1:**
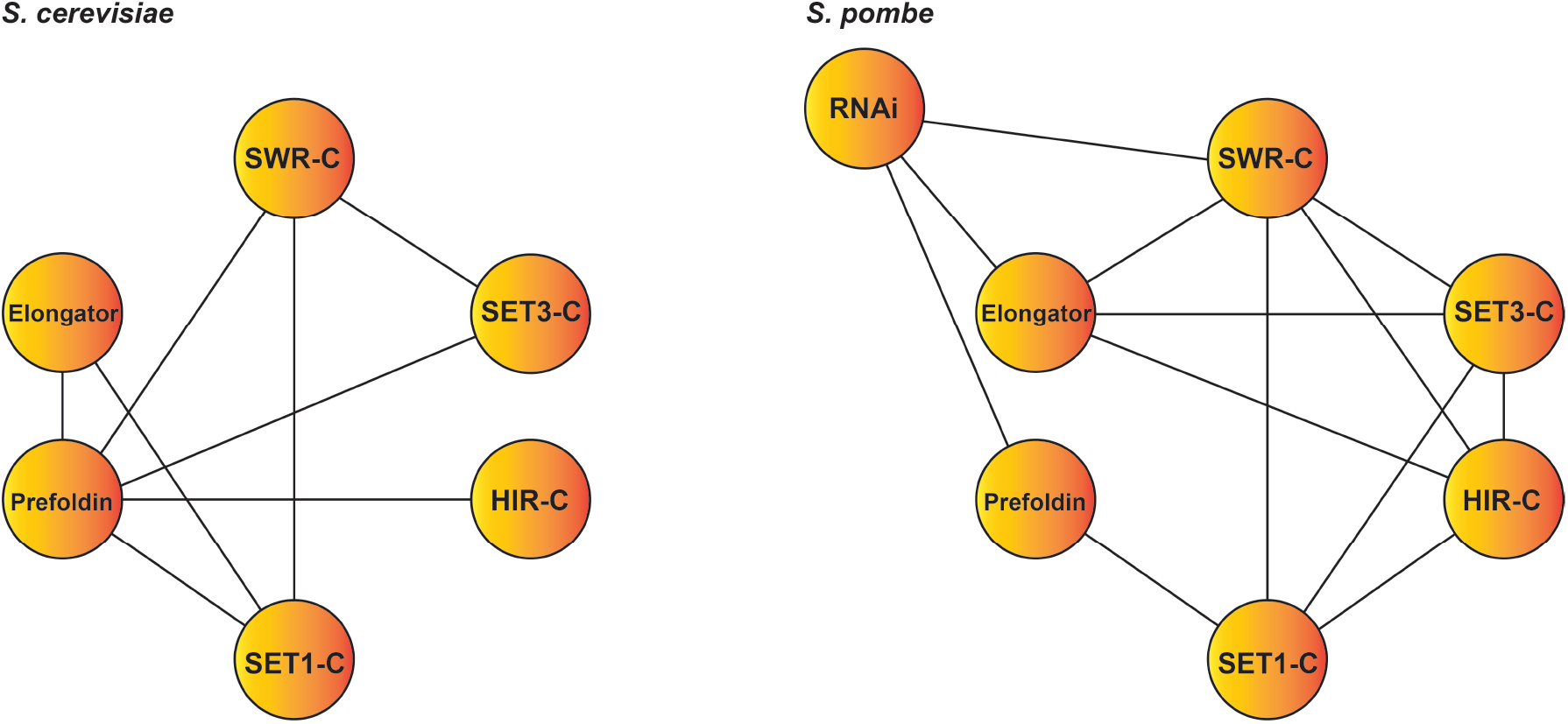
Figure depicting the cross-talk between the functional modules of *Saccharomyces cerevisiae* (Budding yeast) and *Schizosaccharomyces pombe* (Fission yeast) as inferred in ^25^ by exploiting an Epistasis Map. Modules are represented as circles and the interactions between the modules are shown by lines. From this result, the authors of ^25^ conclude that conservation occurs at the level of the functional modules (protein complex), but the cross-talk between the modules differ substantially.

Based on all these above observations, we evidently come to understand that traces of evolution are not only present at the level of the Protein-Protein Interactions, but are also very much present at the level of the inter-module interactions. So, there exists a fair chance of phylogeny reconstruction by analyzing the topology of only the cross-talk between the modules rather than focusing on the conserved intra-module associations. Again, module-based analysis is less time consuming since it mainly relies on the analysis by considering the comprehensive network properties like the network proximity and modularity rather than contemplating the individual interactions of the PPI network ^26^. Therefore, in this chapter, we construct a new network called the Protein Cluster Interaction Network (PCIN) where the spatially localized clusters of the Protein Locality Graph are treated as the nodes and the inter-module associations are considered as the edges. It will be interesting to determine that whether we will be able to reconstruct species trees by exploring the inter-module interactions only. If true, then this will also provide us with an added benefit of reducing the computational burden faced when analyzing the vast Protein Interaction Networks for phylogeny reconstruction. Thus, in this chapter, we propose two hypotheses and perform detailed analyses to check their validations. Firstly, we check whether we are able to decipher biological evolution by exploiting the topology of the Protein Locality Graph. And, secondly, we verify if we are capable of constructing the correct phylogenetic tree of species based solely on the topology of the Protein Cluster Interaction Network, i.e. by focusing on the inter-module connections exclusively. We would like to emphasize here that we do not verify this hypothesis with an underlying objective of proposing a competitive method for the generation of the phylogenetic trees of the biological species; molecular sequence-based existing techniques already address this problem comprehensively. Our main objective is to verify that whether it is possible to generate a correct species tree from the topology of the inter-module connections alone.

### 2.2 Alignment-free network comparison

The most tractable methods for network comparison are those which perform the comparison at the level of the entire network utilizing global features ^27^. But, these global feature-based statistics will never be enough sensitive to be able to shed light on the evolutionary processes ^7^. Hence, local network features are a vital requirement for an efficient comparison of the Protein Interaction Networks. There exist diverse network alignment-based methods which perform network comparisons using the properties of the individual proteins/nodes such as, protein sequence similarity, protein functional similarity, local network similarity, etc ^28–30^. These methods aim to identify matching proteins/nodes between the networks and then use these matching proteins/nodes to recognize the exact or close sub-network matches. There are two primary disadvantages of these methods. Firstly, they are computationally intensive. Secondly, they tend to yield an alignment consisting of only a relatively small proportion of the networks being compared. Though the second disadvantage has been attenuated to some extent by other methods ^31–33^, but still the resulting alignment lacks majority portion of the networks being compared. Again, based on a detailed study conducted on the effect of errors and incompleteness in the alignments of the simulated networks, it has been estimated that only nearly complete networks, possessing > 90% coverage (very rare), have the capability of generating reliable alignments ^34^. Thus, instead of following network alignment paradigm, we perform alignment-free topology-based network comparisons by utilizing the subgraph counts rather than exploiting functional similarity or sequence homology to compare the networks. Network topology refers to the overall layout, or the structure of a network, which means it denotes how different nodes of a network are placed or interconnected with respect to each other. Therefore, alignment-free topology-based methods require only the interaction data for comparing the networks and are conceptually very different from the network alignment-based methods.

### 2.3 Topology-based distance measure

Currently, there is a lack of consensus regarding the likelihood-based statistical models for studying the evolution of the Protein Interaction Networks ^27^. Moreover, distance-based phylogenetic tree reconstruction methods are more robust to mis-specifications and thus, often perform better than the maximum-likelihood methods ^35,36^. Hence, for this study, we utilize Netdis, a one-dimensional alignment-free network comparison statistic which can be exploited to compile a distance matrix for building species trees ^7^. Netdis can successfully reconstruct an accurate species tree for a set of organisms whose significant PPI data is currently available. Another reason for selecting Netdis as our distance measure is its more effective performance as that compared to other alignment-based computationally expensive network comparison statistics like MI-GRAAL ^37^.

Netdis is a topology-based distance measure between networks which is capable of conducting PIN-based phylogeny reconstruction. Since modules and motifs have long been identified as important components of the biological networks and have been already conjectured to play a vital role in evolution by staying conserved among the evolutionary closer species ^5,19–24^, Netdis makes use of the subgraph content to build a distance matrix between the networks. In other words, the primary concept behind Netdis is that similar networks possess similar local neighborhoods. Rather than just counting the total number of subgraphs present in the entire network, Netdis counts the occurrence of subgraph shapes in the local neighborhoods of all the nodes in a network. This is because factors such as network size and density strongly influence the former but not the latter, and thus the former measure becomes too coarse for multiple network comparisons.

Netdis denotes the neighborhood of a protein/node in the Protein Interaction Network by a two-step ego network which is also called as ego-network of radius two ^38^. The two-step ego-network of a protein/node *p* is the (sub) network consisting of all the nodes within two edges of *p*, and also including all the edges between those nodes. For each query network to be compared, Netdis starts by extracting the set of two-step ego-networks for all the nodes of the network. Then, for each two-step ego-network, Netdis counts the number of occurrences of all three to five-node induced subgraphs or graphlets ^22^. A combinatorial subgraph enumeration approach plays the role of the counting algorithm ^39^. Then, a *k*-nodes subgraph count vector (*k* = 3, 4 or 5) gets associated with every node in the network for its corresponding two-step ego-network. These counts are then centred according to the size and the density of the ego-network by utilizing a gold-standard network serving as a proxy for the expected counts. For each *k*-node subgraph in the query network, the centred counts of all the two-step ego-networks are summed up which are then provided as an input to a self-standardizing statistic called *netd*_2_*^S^k*. Finally, Netdis computes a symmetric matrix consisting of the *netd*_2_*^S^k* values for the pair of networks to be compared. Netdis then clusters together the candidate networks by utilizing the resulting symmetric distance matrix and this clustering can be represented by dendograms for further analysis ^7^.

### 2.4 Protein Locality Graph

We consider various pathways from the KEGG pathway maps ^40^ and the PPI information from the interaction databases like Database of Interacting Proteins (DIP) ^41^, Molecular INTeraction Database (MINT) ^42^, IntAct^43^, and BioGRID ^44^ for developing the whole-cell PIN. The PINs are commonly modeled as a graph *G*(*V, E*), where the proteins are denoted as nodes (*V*) and edges (*E*) are the interactions between the pair of proteins. The edges of such networks are undirected and weighted, where the edge weights are used to assimilate reliability information associated to the corresponding PPIs. During the PIN construction, we determined protein locality and allocated the spatial locality scores as the corresponding edge weights. Hence, the constructed whole-cell PIN is also termed as the Protein Locality Graph (PLG).

Markov Cluster Algorithm (MCL) ^45^, an efficient PIN-based clustering method, is used to identify the densely connected sub-graphs/clusters of the PLG which are demonstrated as the functional modules of the whole-cell by a detailed Gene Ontology ^46^ based computational analysis ^17^. A detailed description for constructing the functional modules of the whole-cell can be found at Supplementary Section S1.

### 2.5 Protein Cluster Interaction Network

According to the modular cell biology principles and the speculations mentioned in literature as discussed in the hypothesis (Section 2.1), there exists a fair possibility of finding traces of evolution at the level of the inter-module interactions. So, we may be able to perform phylogeny reconstruction by analyzing the topology of the cross-talk between the protein modules rather than focusing on the conserved intra-module associations. In order to verify this hypothesis, we compress the Protein Locality Graph into a power graph called the Protein Cluster Interaction Network (PCIN).

In computational biology, power graph analysis is a method for the analysis and representation of the complex networks such as, Protein-Protein Interaction Networks, domain-peptide binding motifs, gene regulatory networks, drug-target-disease networks, and social networks. A power graph constitutes a set of power nodes (union of nodes of the original graph) and a set of power edges (edges between power nodes) ^47^. Power graphs are a novel representation of graphs that use cliques, bicliques, and stars as primitives rather than using the ‘node and edge’ language. Power graph analysis can be thought of as a lossless compression algorithm for graphs ^48^ where, compression levels of up to 95% have been obtained for the complex biological networks. Network compression is a new measure derived from the power graphs which has been proposed as a quality measure for the Protein Interaction Networks ^49,50^.

We transform the PLG to the Protein Cluster Interaction Network, a power graph, by denoting the densely connected clusters or functional modules of the PLG as power nodes and the inter-cluster/inter-module connections as power edges, whereas the intra-cluster edges are entirely ignored. While transforming PLG to PCIN, we create a dictionary data structure to store each power node and the subnetwork of the PLG constituting the power node. Thus, PCIN is a completely lossless compressed version of PLG, i.e. no data is lost while converting PLG to PCIN and we can easily restore the exact original state of the PLG from the PCIN if needed. Since in PCIN, the number of PLG nodes gets replaced by the number of clusters, which are very less as compared to the number of PLG nodes (more than 90% reduction happens), therefore, we will achieve a massive reduction in the computational time while performing the topological analysis of the PCIN.

#### Lemma 1.

*Protein Cluster Interaction Network contains all the functional and interaction properties of Protein Locality Graph.*

*Proof.* Modular cell biology states that cellular functions are executed by mod-ules consisting of many species of interacting molecules. According to modular biology, PLGs can be spatially, chemically, and temporally partitioned into functional modules, and can perform biological functions discretely in isolation from the other modules in the network ^15^. Here, PCIN is formed from the functional modules of the PLG, where each functional module becomes a node in PCIN, and the inter-module edges constitute the set of edges in PCIN. The functional modules work discretely in the cell with very less or no functions being performed in collaboration with each other. Hence, PCIN preserves the functional properties of the PLG. When the functional modules of PLG are replaced by corresponding power nodes in PCIN, the subnetworks constituting the modules are stored in a dictionary data structure which can be easily utilized to construct PLG back from PCIN. Thus, PCIN is a complete lossless compressed version of PLG and the interaction properties of PLG are completely preserved during PCIN construction.

### 2.6 Construction of the evolutionary trees

We compute the pairwise Netdis (Section 2.3) values which are then used to build the distance matrix for all the query networks. Unweighted Pair Group Method with Arithmetic mean (UPGMA) ^51^, a simple bottom-up hierarchical clustering method, has been applied over the Netdis distance matrix of the input dataset to generate the final evolutionary/phylogenetic tree. UPGMA is a heuristic greedy method which creates one cluster per network and sequentially merges the nearest pair of clusters by directly using the distance matrix until only two clusters remain. All the trees constructed in this work are rooted bifurcating trees that have exactly two descendants arising from each interior node, and the root corresponds to the most recent common ancestor of all the entities at the leaves of the tree. Here, in this study, we ignore any branch length of the trees since it will be very difficult to interpret them before understanding the effect of errors. We generate the trees using the DendroPy ^52^ package, a library for phylogenetic computing in Python^53^. Here, all the trees have been visualized using an online tool called Interactive Tree Of Life (iTOL) ^54^, which can effectively perform the display, annotation, and management of any phylogenetic tree.

### 2.7 Rand Index

In this study, we determine the accuracy of the resulting dendograms by measuring the similarity in the clusterings by utilizing the Adjusted Rand Index, a variation of the Rand Index. The Rand Index (RI) or the Rand measure is a statistical measure of the similarity between two data clusterings ^55^. RI is related to the accuracy but it is applicable even when the class labels are not used. The Rand index has a value between 0 and 1, with 0 indicating that the two data clusterings do not agree on any pair of points and 1 indicating that the data clusterings are exactly the same. The Adjusted Rand Index (ARI) is the corrected-for-chance version of the Rand Index ^56^. Such a correction for chance establishes a baseline by using the expected similarity of all pair-wise comparisons between clusterings specified by a random model. Though the Rand Index may only yield a value between 0 and +1, the Adjusted Rand Index can yield negative values if the index is less than the expected index. Similar to RI, higher the values of ARI, higher the similarity of the clusterings. Here, we utilize the Adjusted Rand Index to quantify the accuracy of the generated dendograms based on the similarity between the data groupings.

A detailed flow diagram of our proposed methodology is shown in Figure 2.

**Figure 2:**
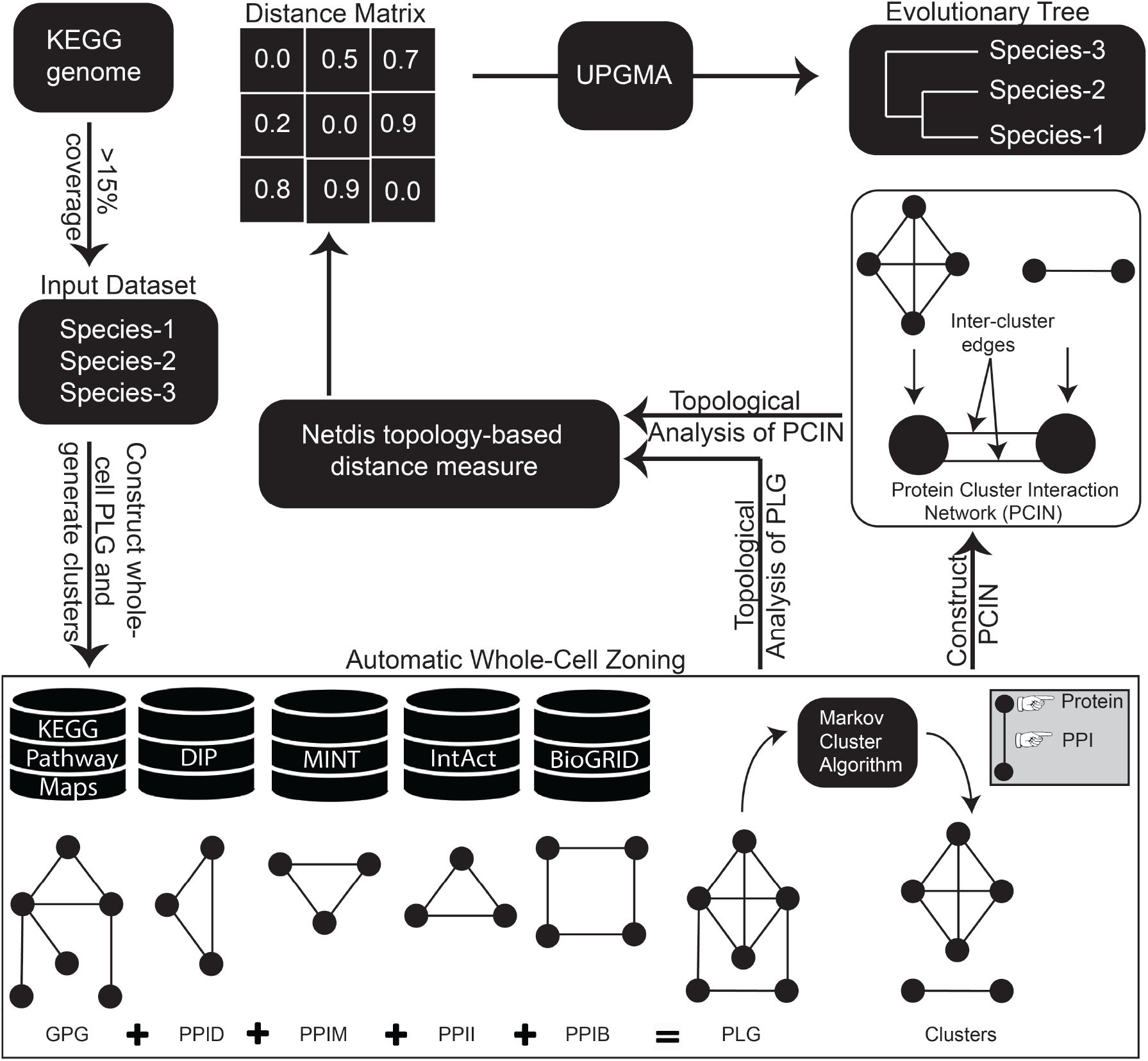
Flow diagram of the Protein Locality Graph and Protein Cluster Interaction Network-based biological evolution.

## 3 Result and Discussions

### 3.1 Simulated networks from random graph models

We initially generate simulated networks from various random graph models in order to perform a statistical analysis of our proposed methodology. Here, we apply the Netdis distance measure to the simulated networks generated from different random graph models and group them together according to the patterns hidden in their topologies. If the different model types differing substantially from each other are correctly grouped together, then we can expect our methodology to correctly cluster together different biological networks also according to their topology-based patterns.

The various random graph models which we consider for this study include — Erdös-Rényi (ER) random graphs ^57^, Watts-Strogatz (WS) model ^58^, Barabási-Albert (BA) model ^59^, and Random Geometric Graph (RGG) model ^60^. We generate the synthetic networks from ER, WS, BA, and RGG random graph models using the Python language package, NetworkX ^61^.

For a given number of nodes and a probability *p* for edge creation, the ER model ^57^ returns a binomial graph also known as the Erdös-Rényi graph. The Erdös-Rényi graph has *n* labeled nodes connected by *m* edges which are randomly chosen from *n*(*n* − 1)/2 possible edges. The Watts-Strogatz random graph model ^58^ accepts three mandatory parameters during the random graph construction. These parameters include *n* number of nodes, *k* number of nearest neighbors in the ring topology to which each node is connected, and the probability *p* of rewiring each edge in the graph. The WS model first creates a ring over *n* nodes. Then each node in the ring is connected with its *k* nearest neighbors. Then some edges are replaced and shortcuts are created, i.e. with probability *p*, each edge *u* − *v* in the underlying ‘*n*-ring with *k* nearest neighbors' is replaced with a new edge *u* − *w* where the node *w* is randomly chosen from the existing *n* nodes. The Barabási-Albert random graph model ^59^ requires two compulsory parameters in order to construct the random graph. These parameters include *n* number of nodes and *m* number of edges to be attached from a new node to the existing nodes. Thus, a Barabási-Albert random graph is grown by adding *m* number of edges to each of the new nodes and these *m* edges are preferentially attached to the existing nodes with high degrees. The Random Geometric Graph model ^60^ requires two necessary parameters while constructing the random graph. These parameters include *n* number of nodes and a threshold value for the radius of the random graph. The RGG model places *n* nodes uniformly at random in the unit cube. Two nodes *u*, *v* are connected with an edge if *d*(*u, v*) ≤ *r*, where *d* is the Euclidean distance and *r* is a radius threshold.

The random networks generated using the ER, WS, BA, and RGG random graph models, play the role of the simulated networks during verification of the presence of traces of evolution in the topology of the Protein Locality Graph. Then, we identify the densely connected clusters of these random networks, utilizing which we transform these random graphs into the power graphs, which we call as Modular Simulated Networks (MSNs), so that they mimic the Protein Cluster Interaction Networks as discussed in Section 2.5. The nature of some of the model generated random networks may not be modular and hence, will not possess a significant tendency to cluster (global clustering coefficient equal to or close to 0). To avoid such cases, we discard any random network with global clustering coefficient < 0.27. Now, in order to generate the clusters, we apply the Markov Cluster Algorithm to the random networks with the inflation parameter of MCL being set dynamically such that no single-node clusters get generated. The power graphs constructed by utilizing the cluster information of the original random networks play the role of the simulated networks during verification of the presence of traces of evolution in the topology of the Protein Cluster Interaction Network where we analyze the inter-module interactions and demonstrate their role in building the species trees.

### 3.2 Protein interaction data

To test our hypothesis on biological data, we consider only those species having at least 500 physical interactions and > 15% coverage ^7^. Here, the term ‘coverage’ denotes a rough estimate of how many proteins have been probed for the interactions given the expected proteome of the organism ^7^. We define the coverage as a percentage measure by dividing the total number of nodes in the Protein Locality Graph by the estimated number of protein coding genes present in the genome of the organism available till date. PLG incorporates the existing experimental PPI data from DIP, MINT, IntAct, and BioGRID. Pathway information from the KEGG Pathway Maps have also been considered which are then mapped to PPIs which are also a part of the PLG. Again, we computationally predict a few PPIs based on spatial locality, which are also added to the PLG. Thus, the PLG consists of a mixture of both experimental as well as predicted protein interaction data. The genes present in the genome of an organism are all reviewed, i.e. manually annotated, whereas, some of the proteins included in the PLG may belong to the unreviewed set and are awaiting full manual annotation as reported by UniProt ^62^. Therefore, the total number of proteins/nodes present in the PLG may sometimes exceed the total estimated number of reviewed genes in the genome of the organism. In total, we analyze five species: *Saccharomyces cerevisiae* (budding yeast), *Arabidopsis thaliana* (thale cress), *Caenorhabditis elegans* (round worm), *Homo sapiens* (human), and *Rattus Norvegicus* (brown rat). Data has been generated for yeast (dated: May 25, 2019), cress (dated: May 27, 2019), worm (dated: May 28, 2019), human (dated: May 09, 2019), and rat (dated: May 09, 2019) by performing real-time web scraping from DIP, MINT, IntAct, BioGRID, and KEGG. Table 1 summarizes the five species' datasets used in this study. We derive the number of protein coding genes present in the genome of the five species from the BioNumbers database ^63^.

**Table 1:** Network summaries for PLG and PCIN data

| Species | Genes | #PLG<br>Nodes | #PLG<br>Edges | PLG<br>coverage | PLG<br>density | #PCIN<br>Nodes | #PCIN<br>Edges | PCIN<br>density |
| --- | --- | --- | --- | --- | --- | --- | --- | --- |
| Yeast | 5616 | 7316 | 544655 | > 100 | 0.02 | 24 | 48 | 0.17 |
| Cress | 20568 | 3473 | 293976 | 16.88 | 0.04 | 116 | 192 | 0.03 |
| Worm | 19735 | 6960 | 81738 | 35.26 | 0.003 | 403 | 1980 | 0.024 |
| Human | 20500 | 41550 | 8943744 | > 100 | 0.01 | 676 | 1660 | 0.007 |
| Rat | 24948 | 9554 | 2652738 | 38.29 | 0.05 | 586 | 7413 | 0.043 |

In all the following results, we set the value of *k* as 4 while applying the Netdis measure ^7^ which associates a *k*-nodes subgraph count vector with every node in the query network during its computation, as mentioned in details in Section 2.3. Though *k* > 4 will surely increase the sensitivity of the method, but an increased computational time requirement for the induced subgraph counting and some preliminary analyses performed in ^7^, render setting *k* > 4 to be futile. Hence, for all the results shown in this section, we utilize Netdis with *k* being set as 4. Again, Netdis computation requires a gold-standard network which serves as a proxy for the expected subgraph counts. Since we test our hypothesis on both simulated and biological networks, we select gold-standard networks from each category. Thus, we select the experimentally verified protein interaction dataset of *Drosophila melanogaster* (fly) from the Database of Interacting Proteins as our gold-standard biological network. Also, we generate a gold-standard simulated network using the Erdös-Rényi model with 4000 nodes and 39992 edges.

### 3.3 Phylogenies from simulated networks

We simulate three networks for each of the four models described in Section 3.1 giving a total of 12 simulated networks. We denote the three out of the 12 simulated networks generated using ER model as *ER*1, *ER*2, and *ER*3. Similarly, we represent the other nine networks generated using WS, BA, and RGG models as, *WS*1, *WS*2, *WS*3, *BA*1, *BA*2, *BA*3, *RGG*1, *RGG*2, and *RGG*3, respectively. Next, we generate the Modular Simulated Networks from each of these 12 random networks following the methodology as mentioned in Section 3.1. We denote the corresponding MSNs derived from *ER*1, *ER*2, *ER*3, *WS*1, *WS*2, *WS*3, *BA*1, *BA*2, *BA*3, *RGG*1, *RGG*2, and *RGG*3, as *M* −*ER*1, *M* −*ER*2, *M* −*ER*3, *M* −*WS*1, *M* −*WS*2, *M* −*WS*3, *M* −*BA*1, *M* −*BA*2, *M* − *BA*3, *M* − *RGG*1, *M* − *RGG*2, and *M* − *RGG*3, respectively.

We perform pairwise alignment-free network comparisons on these 12 query simulated networks and 12 query MSNs separately, and determine their corresponding topology-based distance values using the Netdis measure. We get two pairwise distance matrices for the random networks and their corresponding generated Modular Simulated Networks and the resulting trees are shown in Figure 3. Resulting tree visualizations have been generated using iTOL ^54^. We can see from Figure 3(a) that, the random networks generated using different models are perfectly clustered together according to model type. Also, Figure 3(b) shows that the corresponding Modular Simulated Networks generated from the original random networks are also grouped together correctly according to the model type. Thus, the Adjusted Rand Index of both the dendograms shown in Figure 3(a) and Figure 3(b) is 1.0 indicating that all the samples belong to their correct clusters distinguished by the random graph model categories.

**Figure 3:**
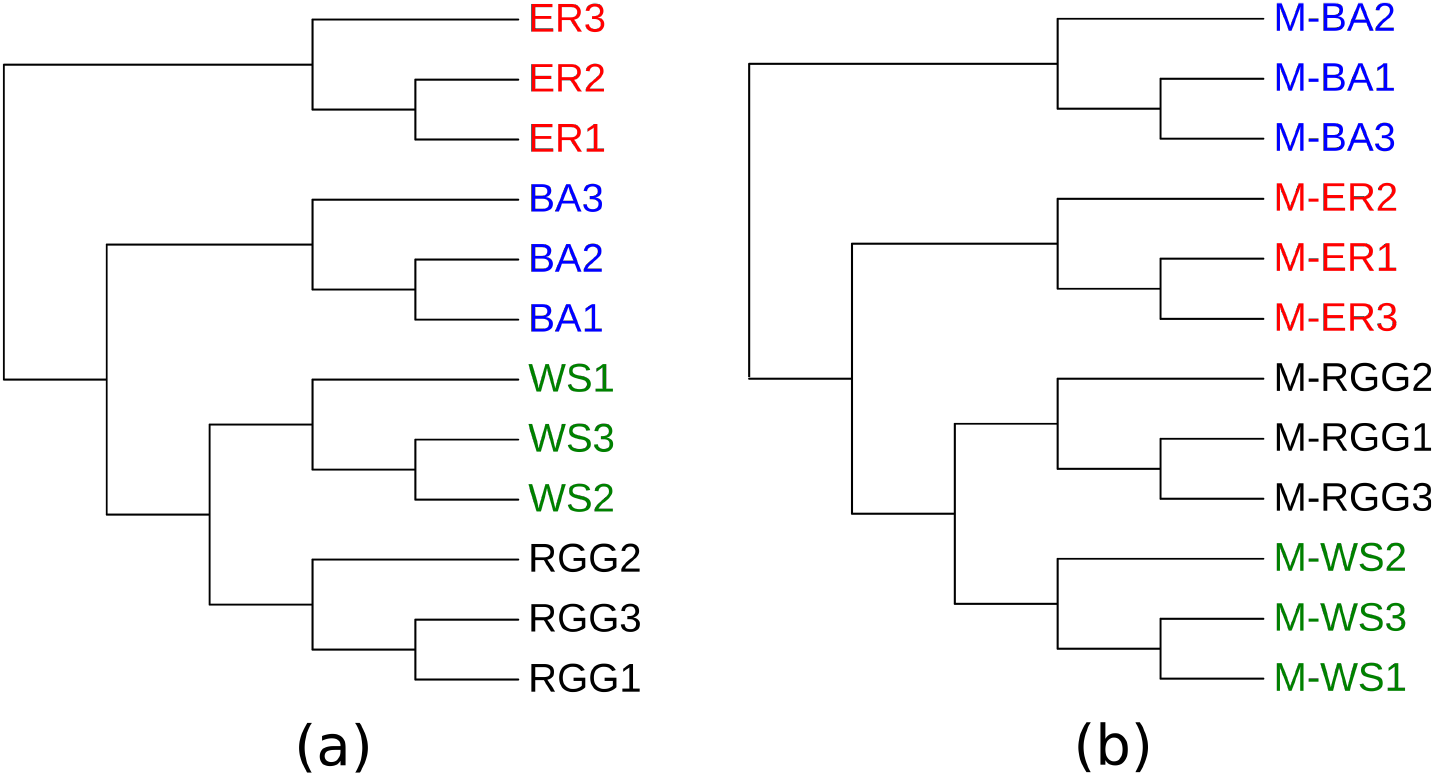
Phylogenetic trees of (a) Simulated Networks having same or similar network size and density. (b) Modular Simulated Networks. Samples from the same model are perfectly clustered together. *ER*1/*ER*2/*ER*3, *WS*1/*WS*2/*WS*3, *BA*1/*BA*2/*BA*3, and *RGG*1/*RGG*2/*RGG*3 denote the simulated networks generated using Erdös-Rényi, Watts-Strogatz, Barabási-Albert, and Random Geometric Graph models respectively. *M* − *ER*1/*M* − *ER*2/*M* − *ER*3, *M* − *WS*1/*M* − *WS*2/*M* − *WS*3, *M* − *BA*1/*M* − *BA*2/*M* − *BA*3, and *M* − *RGG*1/*M* − *RGG*2/*M* − *RGG*3 denote their corresponding Modular Simulated Networks.

In the above analysis, all the original model generated random networks have the same or similar number of nodes and network density. The simulated networks whose dendogram is shown in Figure 3(a) consists of 3000 nodes and similar network density. We also investigate our hypothesis for the random networks possessing varying network size and density. Now, since the number of the generated densely connected clusters vary among the different random networks having same size, so all the MSNs will ultimately have varying network size and density, which is the case for the resulting tree shown in Figure 3(b). This fact is undoubtedly also applicable for those MSNs whose source simulated networks have different network size or density. In order to verify our hypothesis for the simulated networks possessing varying network size or density, we again generate samples using ER, WS, BA, and RGG models with the number of nodes varying between 3000 and 4000. Then, we prepare their corresponding Modular Simulated Networks. Even with such varying sizes, we achieve the correct clustering for both the simulated networks and their corresponding Modular Simulated Networks as shown in Figure 4(a) and Figure 4(b), respectively. Therefore, the ARI measures for both of these dendograms is again 1.0 indicating that all the samples belong to their correct clusters distinguished by the random graph model categories despite the variation in network size and density.

**Figure 4:**
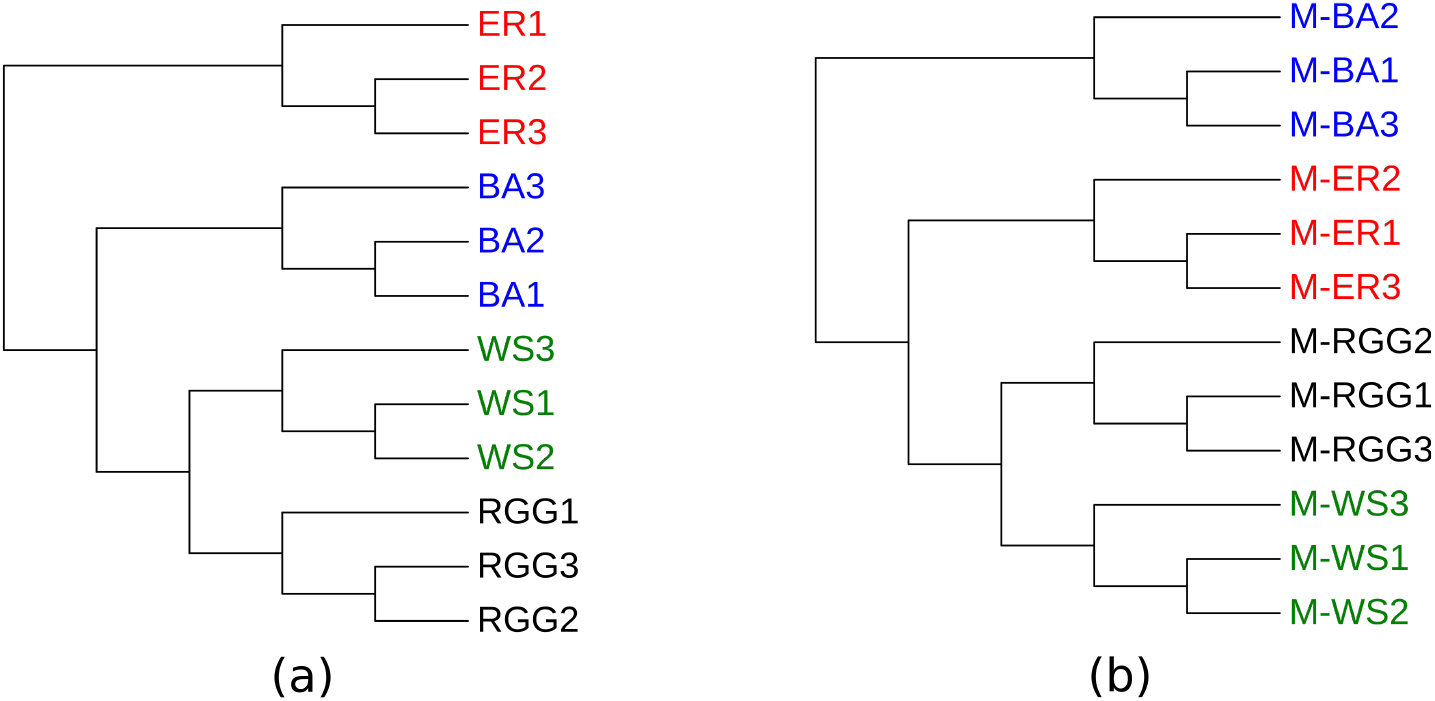
Phylogenetic trees of (a) Simulated Networks of varying network size and density. (b) Modular Simulated Networks. Samples from the same model are perfectly clustered together despite the variation in size and density. *ER*1/*ER*2/*ER*3, *WS*1/*WS*2/*WS*3, *BA*1/*BA*2/*BA*3, and *RGG*1/*RGG*2/*RGG*3 denote the simulated networks generated using Erdös-Rényi, Watts-Strogatz, Barabási-Albert, and Random Geometric Graph models respectively. *M* − *ER*1/*M* − *ER*2/*M* − *ER*3, *M* − *WS*1/*M* − *WS*2/*M* − *WS*3, *M* − *BA*1/*M* − *BA*2/*M* − *BA*3, and *M* − *RGG*1/*M* − *RGG*2/*M* − *RGG*3 denote their corresponding Modular Simulated Networks.

Now, in order to check the robustness of this approach, we perform the above analysis 100 times where each time, we generate 12 sets of simulated networks using ER, WS, BA, RGG models, and their corresponding MSNs. We perform analysis for the fixed node sizes of 1000, 1500, 2000, 2500, 3000, and for variable set of nodes with sizes ranging between 3000 and 4000. We compute the ARI measures of the clusterings generated by applying UPGMA to the Netdis distance matrix. Detailed results are shown in Figure 5 which shows the statistical analysis of the trees generated from the simulated networks of same as well as variable sizes and their corresponding Modular Simulated Networks. The Adjusted Rand Index measures of the generated clusterings are computed with respect to the correct clusterings. From Figure 5, we can observe that, excluding the case where the trees are generated by the simulated networks of 2000 nodes, for all the other cases, the ARI measures are ≥ 0.89 for > 75% of the MSN generated trees in the dataset. Again, except the results for 1000 nodes, from the median lines in the plot we can deduce that ≥ 50% of the MSN generated trees are correct as denoted by their corresponding ARI measures of 1.0. For the simulations performed with 1000 nodes, ≥ 50% of the MSN generated trees have an ARI measure of 0.89. For variable number of nodes, all the random samples in the dataset are correctly grouped together according to the model types and consequently, the ARI measures for all them becomes 1.0 as shown in Figure 5. Apart from these observations, there exist some outliers in the plot (Figure 5) denoting that < 25% of trees in the dataset have those corresponding ARI measures which are marked by the diamonds in the plot for their corresponding simulations. This statistical analysis shows that the random samples from the same model are correctly clustered together, for the majority of the cases, despite the variation in size and density for both the simulated as well as the Modular Simulated Networks. We generate all the results, as shown in Figures 3, 4, and 5, using the Erdös-Rényi model generated simulated network as the gold-standard network required during Netdis computation as mentioned in Sections 3 and 2.3. When we use the DIP dataset of *Drosophila melanogaster* (fly) as the gold-standard network, the resulting trees are same as the ones shown in Figures 3 and 4.

**Figure 5:**
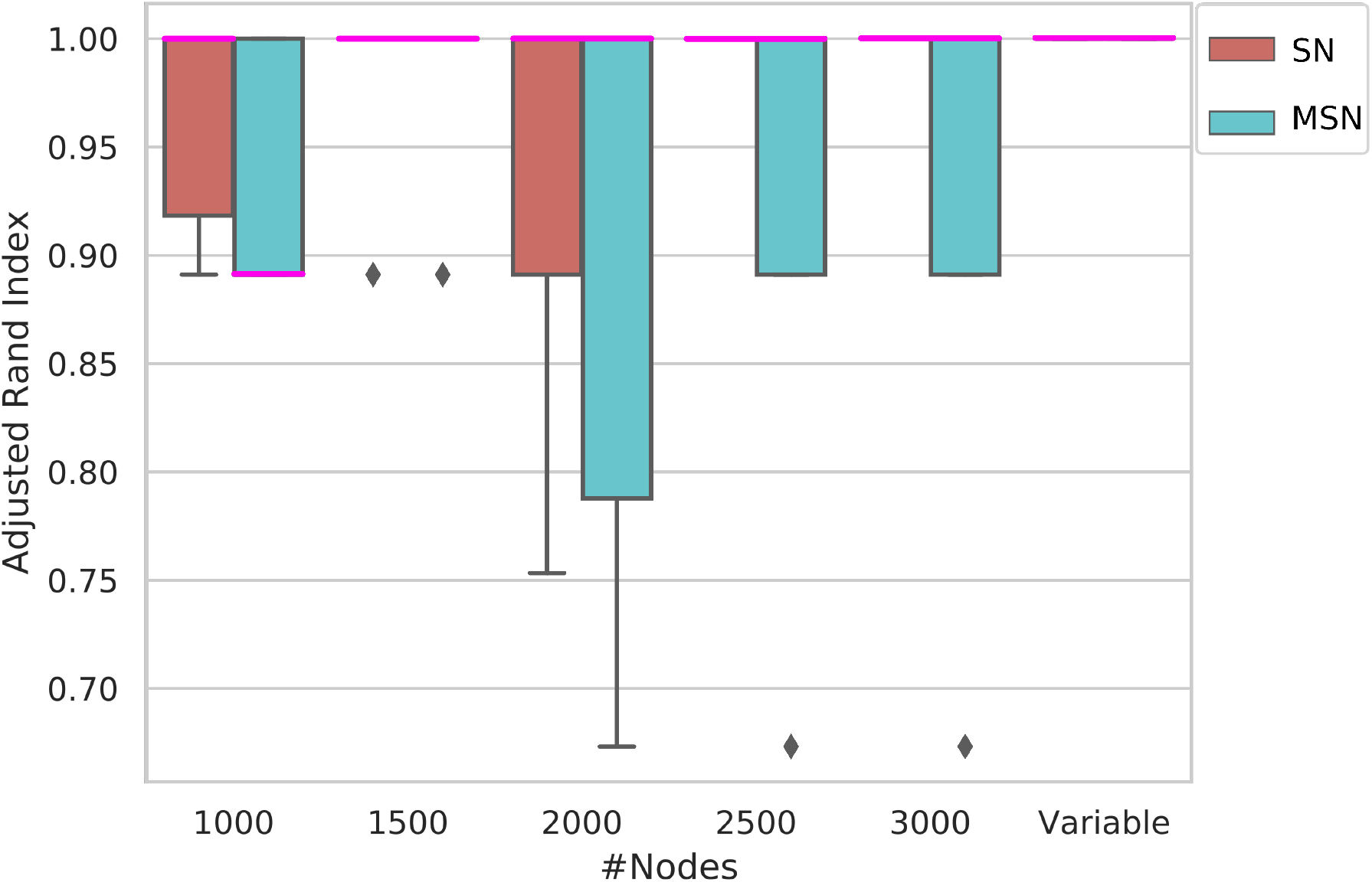
Statistical analysis of the trees generated from the simulated networks of same as well as variable size and their corresponding Modular Simulated Networks. On the Y-axis is the Adjusted Rand Index of similarity with the correct trees or clusterings. Here, SN denotes simulated networks and MSN denotes Modular Simulated Networks. The medians in the plot are marked by pink colored lines. Excluding the case where the trees are generated by the simulated networks of 2000 nodes, for all the other cases, the Adjusted Rand Index measures are ≥ 0.89 for > 75% of the MSN generated trees in the dataset. From the median lines in the plot, we can deduce that for each case except the results for 1000 nodes, ≥ 50% of the MSN generated trees are correct as denoted by their corresponding ARI measures of 1.0. For the simulations performed with 1000 nodes, ≥ 50% of the MSN generated trees have an ARI measure of 0.89. For variable number of nodes, all the random samples in the dataset are correctly grouped together according to the model types. Apart from these observations, there exist some outliers in the plot denoting that < 25% of the trees in the dataset have those corresponding ARI measures which are marked by the diamonds in the plot for their corresponding simulations. For 1500, 2500, 3000 nodes, < 25% of the MSN generated corresponding trees have ARI measure of 0.89, 0.63, and 0.63 respectively. This statistical analysis shows that the random samples from the same model are correctly clustered together, for the majority of the cases, despite the variation in size and density for both the simulated as well as the Modular Simulated Networks.

The simulated networks and the Modular Simulated Networks discussed so far are error-free. Introduction of error will make it harder to differentiate between the networks from the different models. When false positives and false negatives are inserted into the simulated networks of same size and density, then it may be possible to some extent to distinguish the networks according to their model types. However, error has a greater adverse impact when it is already harder to cluster the networks because of their varying sizes and densities. It has been reported in ^7^ that the Netdis measure has a tolerance to an error rate of 50% for the simplest case where error has been introduced to the simulated networks of same size and density. On the other hand, when error has been introduced to the simulated networks of varying sizes and densities, then the Netdis measure display tolerance to an error rate of 5%. However, this tolerance level varies with different datasets which have different sensitivity to error. Thus, when error is present in the Modular Simulated Networks, which are always of varying sizes, Netdis may not be a good measure for our study.

### 3.4 Phylogenies from protein interaction data

The currently accepted phylogeny between the species listed in Table 1 is represented by the tree in Figure 6(a). We extract this tree from the NCBI taxonomy database ^64^ which incorporates different phylogenetic resources for generating the trees. The phylogenetic tree of our constructed Protein Cluster Interaction Network, which is the modular network generated from the Protein Locality Graph, generated using Netdis is shown in Figure 6(b). Also, the tree generated from the PLG is the same as that shown in Figure 6(b). Out of the multiple possible rooted trees with five leaves, we obtain the correct clustering with human next to rat in one cluster, and the worm, cress, and yeast in separate clusters. For PLG, we get this result despite the notable differences in their node size, edge size, network density, and coverage as mentioned in Table 1. For PCIN, we get this result despite a significant compression of the PLG and reduction in the network sizes (Table 1). Netdis possesses a tolerance to an error rate of 50% in case of the Protein Interaction Networks ^7^. So, there exists a fairly good possibility for Netdis to perform a correct alignment-free comparison of networks bearing false positives and false negatives. Thus, although PLG consists of protein spatial locality-based predicted PPIs (which may be false positive), we obtain the correct clustering as shown by the result in Figure 6(b). Surprisingly, for the Protein Cluster Interaction Network, a novel modular network constructed from the Protein Locality Graph by preserving the inter-module connections and ignoring the intra-module connections, we get the correct clustering as depicted by Figure 6. An added benefit of utilizing PCIN is the very less computational time requirement during its topological analysis as that compared to the PINs or the PLG. Therefore, we can conclude that, in spite of the lack of conservation of individual PPIs, the topology of the Protein Locality Graph contains relevant information about the evolutionary processes. Also, from the detailed PCIN topology-based analysis, we can say that evolution tinkers mainly with the inter-cluster/inter-module connections and the intra-module connections play little to no role in the evolutionary processes. Therefore, to summarize, we claim our proposed hypotheses, as mentioned in Section 2.1, to be true.

**Figure 6:**
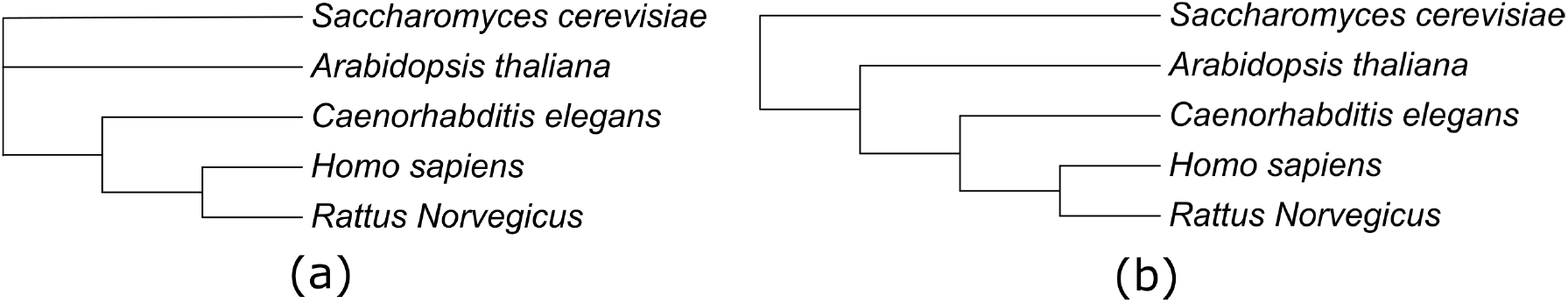
Phylogenetic trees from (a) NCBI taxonomy database^64^. (b) Protein Cluster Interaction Network. Topology-based analysis of the Protein Locality Graph also generates the same tree as Protein Cluster Interaction Network as shown in (b).

## 4 Conclusions

In this paper, we provide evidence for the hypothesis that the topology of the Protein Interaction Networks alone contain evolutionary information without any supplementary biological data. Our results reveal that the currently available PPI data are sufficiently profuse for deriving the correct evolutionary relationships, at least between the model species. We establish that evolutionary relationships among the species, having PPI data with the genome coverage of at least 15%, can be discovered following a detailed topological analysis of the Protein Locality Graph as well as module-based topological analysis of the Protein Cluster Interaction Network. We construct the PLG by collecting PPI data from several interaction databases and the pathway maps based on an automatic whole-cell zoning method, whereas, the PCIN is a modular compressed form of the PLG. We utilize Netdis measure for perforing their alignment-free network comparisons and generating the distance matrices for constructing UPGMA-based evolutionary trees.

We emphasize that we do not propose our contribution as a competitive method for generating phylogenetic trees of the species and the existing molecular sequence-based techniques address this problem effectively. Our main result is that we can derive a correct phylogenetic tree from the topology of Protein Locality Graph, a spatial locality-based Protein Interaction Network. Also, we can get the species trees by focusing on the inter-protein-module connections only. We find that evolution tinkers mainly with the inter-cluster/inter-module connections and the intra-module connections play little to no role in the evolutionary processes, a claim also made in literature ^5,15,19–24^. The biological perspective of this work is that closely-related species will on average share more topologically similar PIN neighborhoods than the unrelated species. Again, a biological function is carried out in a cell by communities of interacting proteins and closely-related species will have on average more of these communities in common ^7^. Since PCIN has been constructed based on the Modular Biology principles, deriving evolutionary relationships based on the topological analysis of PCIN opens up a new branch of research viz., Evolutionary Modular Biology. More studies should be conducted on the evolution of the Protein Interaction Networks and the inter-module connections, which will shed light on the evolution of species with time from a biological networks' perspective. These contributions will nevertheless be a major progressive step in the domain of the Evolutionary Systems Biology.

## References

[1] M. W. Gray, G. Burger, and B. F. Lang, “Mitochondrial Evolution,” Science, vol. 283, no. 5407, pp. 1476–1481, 1999.

[2] M. Hegreness and R. Kishony, “Analysis of genetic systems using experimental evolution and whole-genome sequencing,” Genome Biology, vol. 8, no. 1, pp. 1–5, 2007.

[3] D. M. Krylov, Y. I. Wolf, I. B. Rogozin, and E. V. Koonin, “Gene Loss, Protein Sequence Divergence, Gene Dispensability, Expression Level, and Interactivity Are Correlated in Eukaryotic Evolution,” Genome Research, vol. 13, no. 10, pp. 2229–2235, 2003.

[4] M. Yokono, S. Satoh, and A. Tanaka, “Comparative analyses of whole-genome protein sequences from multiple organisms,” Scientific Reports, vol. 8, no. 1, pp. 1–10, 2018.

[5] S. Erten, X. Li, G. Bebek, J. Li, and M. Koyutürk, “Phylogenetic analysis of modularity in protein interaction networks,” BMC Bioinformatics, vol. 10, no. 1, p. 333, 2009.

[6] Z. Liang, M. Xu, M. Teng, and L. Niu, “Comparison of protein interaction networks reveals species conservation and divergence,” BMC Bioinformatics, vol. 7, no. 1, p. 457, 2006.

[7] W. Ali, T. Rito, G. Reinert, F. Sun, and C. M. Deane, “Alignment-free protein interaction network comparison,” Bioinformatics, vol. 30, no. 17, pp. i430–i437, 2014.

[8] Y. Jin, D. Turaev, T. Weinmaier, T. Rattei, and H. A. Makse, “The Evolutionary Dynamics of Protein-Protein Interaction Networks Inferred from the Reconstruction of Ancient Networks,” PloS One, vol. 8, no. 3, p. e58134, 2013.

[9] Q. Zhong, S. J. Pevzner, T. Hao, Y. Wang, R. Mosca, J. Menche, M. Taipale, M. Taşan, C. Fan, X. Yang, et al., “An inter-species protein–protein interaction network across vast evolutionary distance,” Molecular systems biology, vol. 12, no. 4, p. 865, 2016.

[10] R. Sharan, S. Suthram, R. M. Kelley, T. Kuhn, S. McCuine, P. Uetz, T. Sittler, R. M. Karp, and T. Ideker, “Conserved patterns of protein interaction in multiple species,” Proceedings of the National Academy of Sciences of the United States of America, vol. 102, no. 6, pp. 1974–1979, 2005.

[11] A.-L. Barabasi and Z. N. Oltvai, “Network biology: understanding the cell's functional organization,” Nature Reviews Genetics, vol. 5, no. 2, pp. 101–113, 2004.

[12] B. Chor and T. Tuller, “Biological Networks: Comparison, Conservation, and Evolution via Relative Description Length,” Journal of Computational Biology, vol. 14, no. 6, pp. 817–838, 2007.

[13] F. Jordán, T.-P. Nguyen, and W.-c. Liu, “Studying protein–protein interaction networks: a systems view on diseases,” Briefings in Functional Genomics, vol. 11, no. 6, pp. 497–504, 2012.

[14] R. Sharan and T. Ideker, “Modeling cellular machinery through biological network comparison,” Nature Biotechnology, vol. 24, no. 4, pp. 427–433, 2006.

[15] L. H. Hartwell, J. J. Hopfield, S. Leibler, and A. W. Murray, “From molecular to modular cell biology,” Nature, vol. 402, no. 6761supp, p. C47, 1999.

[16] D. Waxman and J. R. Peck, “Pleiotropy and the Preservation of Perfection,” Science, vol. 279, no. 5354, pp. 1210–1213, 1998.

[17] B. Das, A. R. Patil, and P. Mitra, “A network-based zoning for parallel whole-cell simulation,” Bioinformatics, vol. 35, no. 1, pp. 88–94, 2019.

[18] A. Schoenrock, D. Burnside, H. Moteshareie, S. Pitre, M. Hooshyar, J. R. Green, A. Golshani, F. Dehne, and A. Wong, “Evolution of protein-protein interaction networks in yeast,” PLoS One, vol. 12, no. 3, p. e0171920, 2017.

[19] G. P. Wagner, M. Pavlicev, and J. M. Cheverud, “The road to modularity,” Nature Reviews Genetics, vol. 8, no. 12, pp. 921–931, 2007.

[20] X. Zhu, M. Gerstein, and M. Snyder, “Getting connected: analysis and principles of biological networks,” Genes & Development, vol. 21, no. 9, pp. 1010–1024, 2007.

[21] A. P. Cootes, S. H. Muggleton, and M. J. Sternberg, “The Identification of Similarities between Biological Networks: Application to the Metabolome and Interactome,” Journal of Molecular Biology, vol. 369, no. 4, pp. 1126–1139, 2007.

[22] N. Pržulj, “Biological network comparison using graphlet degree distribution,” Bioinformatics, vol. 23, no. 2, pp. e177–e183, 2007.

[23] B. Snel and M. A. Huynen, “Quantifying Modularity in the Evolution of Biomolecular Systems,” Genome Research, vol. 14, no. 3, pp. 391–397, 2004.

[24] A. Vespignani, “Evolution thinks modular,” Nature Genetics, vol. 35, no. 2, pp. 118–119, 2003.

[25] A. Roguev, S. Bandyopadhyay, M. Zofall, K. Zhang, T. Fischer, S. R. Collins, H. Qu, M. Shales, H.-O. Park, J. Hayles, et al., “Conservation and Rewiring of Functional Modules Revealed by anEpistasis Map in Fission Yeast,” Science, vol. 322, no. 5900, pp. 405–410, 2008.

[26] B. Titz, M. Schlesner, and P. Uetz, “What do we learn from high-throughput protein interaction data?,” Expert Review of Proteomics, vol. 1, no. 1, pp. 111–121, 2004.

[27] O. Ratmann, C. Wiuf, and J. W. Pinney, “From evidence to inference: Probing the evolution of protein interaction networks,” HFSP Journal, vol. 3, no. 5, pp. 290–306, 2009.

[28] J. Flannick, A. Novak, C. B. Do, B. S. Srinivasan, and S. Batzoglou, “Automatic Parameter Learning for Multiple Network Alignment,” in Annual International Conference on Research in Computational Molecular Biology, pp. 214–231, Springer, 2008.

[29] H. T. Phan and M. J. Sternberg, “PINALOG: a novel approach to align protein interaction networks—implications for complex detection and function prediction,” Bioinformatics, vol. 28, no. 9, pp. 1239–1245, 2012.

[30] R. Singh, J. Xu, and B. Berger, “Global alignment of multiple protein interaction networks with application to functional orthology detection,” Proceedings of the National Academy of Sciences of the United States of America, vol. 105, no. 35, pp. 12763–12768, 2008.

[31] F. Alkan and C. Erten, “BEAMS: backbone extraction and merge strategy for the global many-to-many alignment of multiple PPI networks,” Bioinformatics, vol. 30, no. 4, pp. 531–539, 2014.

[32] J. Hu, B. Kehr, and K. Reinert, “NetCoffee: a fast and accurate global alignment approach to identify functionally conserved proteins in multiple networks,” Bioinformatics, vol. 30, no. 4, pp. 540–548, 2014.

[33] R. Patro and C. Kingsford, “Global network alignment using multiscale spectral signatures,” Bioinformatics, vol. 28, no. 23, pp. 3105–3114, 2012.

[34] W. Ali and C. M. Deane, “Evolutionary analysis reveals low coverage as the major challenge for protein interaction network alignment,” Molecular BioSystems, vol. 6, no. 11, pp. 2296–2304, 2010.

[35] G. H. Gonnet, “Surprising results on phylogenetic tree building methods based on molecular sequences,” BMC Bioinformatics, vol. 13, no. 1, p. 148, 2012.

[36] J. P. Huelsenbeck and D. M. Hillis, “Success of Phylogenetic Methods in the Four-Taxon Case,” Systematic Biology, vol. 42, no. 3, pp. 247–264, 1993.

[37] O. Kuchaiev and N. Pržulj, “Integrative network alignment reveals large regions of global network similarity in yeast and human,” Bioinformatics, vol. 27, no. 10, pp. 1390–1396, 2011.

[38] P. Pattison, Algebraic Models for Social Networks, vol. 7. Cambridge University Press, 1993.

[39] T. Hočevar and J. Demšar, “A combinatorial approach to graphlet counting,” Bioinformatics, vol. 30, no. 4, pp. 559–565, 2014.

[40] M. Kanehisa, M. Furumichi, M. Tanabe, Y. Sato, and K. Morishima, “KEGG: new perspectives on genomes, pathways, diseases and drugs,” Nucleic Acids Research, vol. 45, no. D1, pp. D353–D361, 2016.

[41] L. Salwinski, C. S. Miller, A. J. Smith, F. K. Pettit, J. U. Bowie, and D. Eisenberg, “The Database of Interacting Proteins: 2004 update,” Nucleic Acids Research, vol. 32, no. suppl 1, pp. D449–D451, 2004.

[42] L. Licata, L. Briganti, D. Peluso, L. Perfetto, M. Iannuccelli, E. Galeota, F. Sacco, A. Palma, A. P. Nardozza, E. Santonico, et al., “MINT, the molecular interaction database: 2012 update,” Nucleic Acids Research, vol. 40, no. D1, pp. D857–D861, 2011.

[43] S. Kerrien, B. Aranda, L. Breuza, A. Bridge, F. Broackes-Carter, C. Chen, M. Duesbury, M. Dumousseau, M. Feuermann, U. Hinz, et al., “The IntAct molecular interaction database in 2012,” Nucleic Acids Research, vol. 40, no. D1, pp. D841–D846, 2011.

[44] A. Chatr-Aryamontri, R. Oughtred, L. Boucher, J. Rust, C. Chang, N. K. Kolas, L. O’Donnell, S. Oster, C. Theesfeld, A. Sellam, et al., “The BioGRID interaction database: 2017 update,” Nucleic Acids Research, vol. 45, no. D1, pp. D369–D379, 2017.

[45] S. Dongen, “A cluster algorithm for graphs,” 2000.

[46] G. O. Consortium, “The Gene Ontology Resource: 20 years and still GOing strong,” Nucleic Acids Research, vol. 47, no. D1, pp. D330–D338, 2019.

[47] S. E. Ahnert, “Generalised power graph compression reveals dominant relationship patterns in complex networks,” Scientific Reports, vol. 4, p. 4385, 2014.

[48] M. Dehmer, F. Emmert-Streib, and S. Pickl, Computational Network Theory: Theoretical Foundations and Applications. John Wiley & Sons, 2015.

[49] L. Royer, M. Reimann, A. F. Stewart, and M. Schroeder, “Network Compression as a Quality Measure for Protein Interaction Networks,” PloS One, vol. 7, no. 6, p. e35729, 2012.

[50] L. Royer, M. Reimann, B. Andreopoulos, and M. Schroeder, “Unraveling Protein Networks with Power Graph Analysis,” PLoS Computational Biology, vol. 4, no. 7, p. e1000108, 2008.

[51] R. R. Sokal, “A statistical method for evaluating systematic relationships,” Univ. Kansas, Sci. Bull., vol. 28, pp. 1409–1438, 1958.

[52] J. Sukumaran and M. T. Holder, “DendroPy: a Python library for phylogenetic computing,” Bioinformatics, vol. 26, no. 12, pp. 1569–1571, 2010.

[53] T. E. Oliphant, “Python for Scientific Computing,” Computing in Science & Engineering, vol. 9, no. 3, pp. 10–20, 2007.

[54] I. Letunic and P. Bork, “Interactive Tree Of Life (iTOL) v4: recent updates and new developments,” Nucleic Acids Research, vol. 47, no. W1, pp. W256–W259, 2019.

[55] W. M. Rand, “Objective Criteria for the Evaluation of Clustering Methods,” Journal of the American Statistical Association, vol. 66, no. 336, pp. 846–850, 1971.

[56] L. Hubert and P. Arabie, “Comparing partitions,” Journal of Classification, vol. 2, no. 1, pp. 193–218, 1985.

[57] P. Erdös and A. Rényi, “On the evolution of random graphs,” Publ. Math. Inst. Hung. Acad. Sci, vol. 5, no. 1, pp. 17–60, 1960.

[58] D. J. Watts and S. H. Strogatz, “Collective dynamics of 'small-world’ networks,” Nature, vol. 393, no. 6684, pp. 440–442, 1998.

[59] A.-L. Barabási and R. Albert, “Emergence of Scaling in Random Networks,” Science, vol. 286, no. 5439, pp. 509–512, 1999.

[60] M. Penrose, Random Geometric Graphs, vol. 5. Oxford university press, 2003.

[61] A. Hagberg, P. Swart, and D. S Chult, “Exploring Network Structure, Dynamics, and Function using NetworkX,” in Proceedings of the 7th Python in Science Conference (SciPy 2008), 2008.

[62] T. U. Consortium, “The Universal Protein Resource (UniProt) in 2010,” Nucleic Acids Research, vol. 38, no. suppl 1, pp. D142–D148, 2009.

[63] R. Milo, P. Jorgensen, U. Moran, G. Weber, and M. Springer, “BioNumbers—the database of key numbers in molecular and cell biology,” Nucleic Acids Research, vol. 38, no. suppl 1, pp. D750–D753, 2010.

[64] S. Federhen, “The NCBI Taxonomy database,” Nucleic Acids Research, vol. 40, no. D1, pp. D136–D143, 2012.

